# Single cell resolution of SARS-CoV-2 tropism, antiviral responses, and susceptibility to therapies in primary human airway epithelium

**DOI:** 10.1101/2020.10.19.343954

**Authors:** Jessica K. Fiege, Joshua M. Thiede, Hezkiel Nanda, William E. Matchett, Patrick J. Moore, Noe Rico Montanari, Beth K. Thielen, Jerry Daniel, Emma Stanley, Ryan C. Hunter, Vineet D. Menachery, Steven S. Shen, Tyler D. Bold, Ryan A. Langlois

**Author notes:** Authors contributed equally (Jessica K. Fiege, Joshua M. Thiede, Hezkiel Nanda, Steven S. Shen, Tyler D. Bold, Ryan A. Langlois).

## Abstract

The human airway epithelium is the initial site of SARS-CoV-2 infection. We used flow cytometry and single cell RNA-sequencing to understand how the heterogeneity of this diverse cell population contributes to elements of viral tropism and pathogenesis, antiviral immunity, and treatment response to remdesivir. We found that, while a variety of epithelial cell types are susceptible to infection, ciliated cells are the predominant cell target of SARS-CoV-2. The host protease TMPRSS2 was required for infection of these cells. Importantly, remdesivir treatment effectively inhibited viral replication across cell types, and blunted hyperinflammatory responses. Induction of interferon responses within infected cells was rare and there was significant heterogeneity in the antiviral gene signatures, varying with the burden of infection in each cell. We also found that heavily infected secretory cells expressed abundant IL-6, a potential mediator of COVID-19 pathogenesis.

## Introduction

SARS-CoV-2, the virus responsible for COVID-19, primarily infects cells of the respiratory tract. The cellular tropism of SARS-CoV-2 may impact several aspects of the disease, including viral spread within and between hosts, mechanisms of immune control of infection or tissue pathology, and the therapeutic response to promising antivirals. Normal human tracheal bronchial epithelial (nHTBE) cells represent a diverse mix of ciliated epithelial cells, secretory cells, and basal cells that form a pseudostratified epithelium when cultured at the air-liquid interface, phenocopying the upper airway in humans [1, 2]. Importantly, cells in this culture system also express endogenous levels of critical host factors including ACE2 and host proteases such as TMPRSS2 that are needed for SARS-CoV-2 viral entry [3-7]. This model also demonstrates key aspects of host antiviral epithelial immunity [8, 9]. Recently, several studies using primary human lung cell cultures and respiratory cells isolated from SARS-CoV-2 infected patients have identified SARS-CoV-2 tropism for ciliated and secretory cells in the upper airway [10-13]. However, the heterogeneity of virus replication and induction of antiviral genes and proinflammatory cytokines within these cells is still unknown.

Remdesivir (GS-5734) has emerged as a promising direct antiviral therapy against SARS-CoV-2, with potent activity demonstrated against several coronaviruses [14, 15]. A landmark clinical trial found that remdesivir treatment of hospitalized individuals with COVID-19 improved median recovery time [16], and this drug is now approved for COVID-19 under emergency use authorization by the U.S. Food and Drug Administration. Remdesivir is a prodrug that is metabolized in cells to the nucleotide analog remdesivir triphosphate, which interferes with coronavirus replication through delayed RNA chain termination [10, 17-19]. Recent studies have identified differential efficacy of remdesivir against SARS-CoV-2 in a range of cell culture systems linked to metabolism of the prodrug to the active form [10]. In addition to differential metabolism, other factors that may impact the variable efficacy of this drug in different cell types include differential drug uptake and heterogeneous permissibility of each cell type to viral entry and replication. While remdesivir clearly exhibits antiviral activity against SARS-CoV-2 in nHTBE cultures, it is not known if there are cell type-dependent differences in drug efficacy.

Following infection coronaviruses are recognized by MDA5 and RIGI leading to the production of type I and III interferons (IFNs), which induce transcriptional programs that mobilize cellular antiviral defenses. Coronaviruses use several mechanisms to successfully evade detection resulting in rare and heterogeneous IFN production [20, 21], similar to influenza virus infected cells [22-25]. During influenza A virus infection, we have previously identified interferon stimulated genes (ISGs) specifically induced in cells supporting high levels of virus replication and we have defined cell type-specific ISGs [26, 27]. Additionally, we and others have found significant heterogeneity in antiviral responses across different cell types [27, 28]. Cell type-specific responses and the degree of heterogeneity in antiviral responses can dictate the outcome of immune responses and infection.

Here, we use nHTBE cells infected with SARS-CoV-2 to demonstrate that remdesivir reduces viral replication uniformly in all susceptible cell types within the upper respiratory tract. Additionally, we demonstrate that TMPRSS2 is the primary host protease used for SARS-CoV-2 entry across cell types in the upper airway. Using single cell RNA sequencing, we further define SARS-CoV-2 tropism and the induction of antiviral and proinflammatory immune responses. We also uncover cellular heterogeneity in SARS-CoV-2 replication within the human airway epithelium and determine the relationship between productive viral infection and induction of IFN. Additionally, we define respiratory epithelial ISG expression in response to SARS-CoV-2 infection and uncover epithelial cell type-specific ISGs, and ISGs associated with either high levels of viral replication or protection from infection. Finally, we demonstrate that the proinflammatory cytokine IL-6 is primarily upregulated within heavily infected secretory cells. Together, these data help to define the early SARS-CoV-2 replication and antiviral landscape within the respiratory tract.

## Results

### SARS-CoV-2 tropism and cell type-specific efficacy of remdesivir in differentiated tracheal bronchial epithelial cells

To determine how SARS-CoV-2 tropism varies within the upper airway we infected nHTBE cells differentiated at air-liquid interface with a SARS-CoV-2 infectious clone generated to express the fluorescent protein mNeonGreen (mNG) [29]. Using flow cytometry, we detected 12.5% of untreated nHTBE cells expressing mNG after icSARS-CoV-2-mNG infection (Fig. 1 A and B); treatment with remdesivir or IFNβ substantially reduced the percentage of mNG positive cells showing clear antiviral effect. We subsequently identified different cell types using the expression of several surface markers and SiR-tubulin (Fig. 1C) and found that the ratio of different cell types in each culture remained similar during infection, regardless of anti-viral treatments (Fig. 1D). Among mNG^+^ SARS-CoV-2 infected cells, we discovered a preponderance of ciliated (SiR-tubulin^+^ CD271^-^) and few infected secretory cells (CD66c^+^ SiR-tubulin^-^ CD271^-^) (Fig. 1E). We also identified a subset of infected cells lacking cell type-specific markers (termed “triple negative”) and a population of cells expressing intermediate levels of CD271 and CD66c, but negative for SiR-tubulin, termed “other”. We did not identify mNG+ basal cells (CD271^+^ SiR-tubulin^-^ CD66c^-^). We also observed that ciliated cells expressed the highest mNG MFI, suggesting that this cell type is the most permissive for SARS-CoV-2 replication (Fig. 1F). These data are consistent with recent reports demonstrating ciliated cells as a dominant target cell of SARS-CoV-2 [10-13, 30], and they suggest that basal cells are relatively resistant to infection. We next evaluated if the antiviral activity of remdesivir varies in different cell types depending on cellular uptake and metabolism [10], which could differ dramatically within the diverse population of cells in the upper respiratory tract. To determine how remdesivir impacts viral replication across all susceptible cells in the upper airway, we pretreated nHTBE cells with remdesivir at 0.1μM, a dose expected to inhibit 80% of SARS-CoV-2 replication [10] and infected with icSARS-CoV-2-mNG. As a positive control for antiviral activity, we also pretreated cells with IFNβ, which completely abrogated infection in nHTBE cells (Fig. 1A and B), consistent with recent results demonstrating sensitivity of SARS-CoV-2 to interferon [31-33]. Remdesivir reduced both the number of SARS-CoV-2 infected cells by ∼5-fold and the expression level of mNG within infected cells (Fig. 1B and F). Importantly, among infected cells, we found that remdesivir similarly inhibited replication regardless of cell type (Fig. 1E). These data suggest that cell type heterogeneity within the respiratory tract does not impact remdesivir efficacy.

**Fig. 1.**
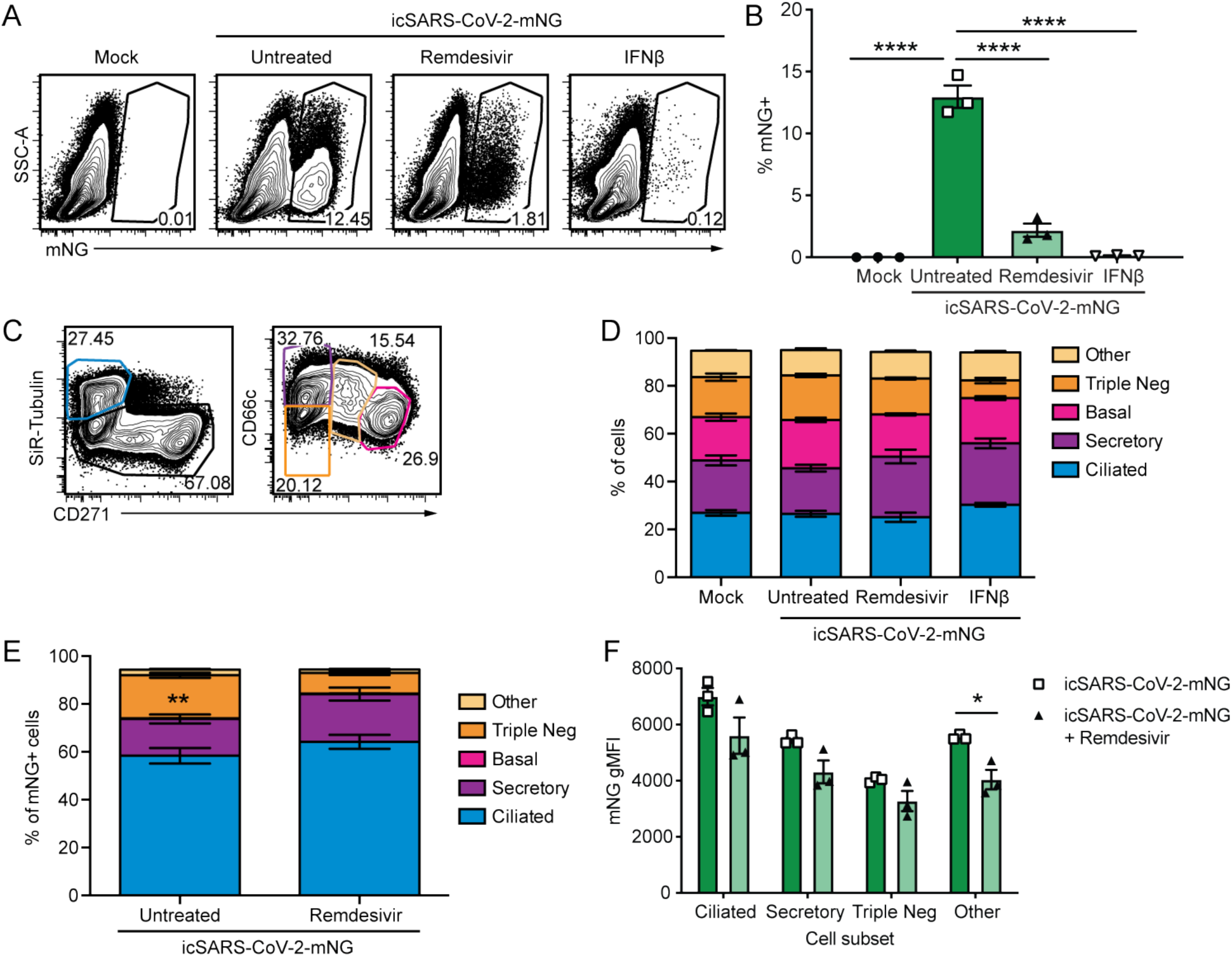
SARS-CoV-2 infects multiple cell types and infection is controlled by remdesivir in primary nHTBE cells. nHTBE cells differentiated at air-liquid interface were untreated or treated with IFN-β or remdesivir prior to infection with icSARS-CoV-2-mNG at a MOI of 2.5. Cells were analyzed 48 hpi by flow cytometry. (A) Representative plots of live cells (B) Frequency of infected cells (mNG^+^). (C) Representative plots of cell subset gating. Ciliated (SiR-Tubulin^+^ CD271^-^), secretory (SiR-Tubulin^-^ CD271^-^ CD66c^+^), basal (SiR-Tubulin^-^ CD271^+^ CD66c^-^), Triple negative (SiR-Tubulin^-^ CD271^-^ CD66c^-^) and other cells (SiR-Tubulin^-^ CD217^int^ CD66c^int^) (D) Frequency of cell subsets from live cells. (E) Frequency of cell subsets of infected cells (mNG^+^). (F) mNG gMFI within each cell subset. The data (B, D-F) are 3 biological replicates per group +/-SEM, 1 of 2 independent experiments.

### Single cell RNA sequencing reveals heterogeneity in SARS-CoV-2 infection across differentiated tracheal bronchial epithelial cells

Tracheal-bronchial epithelial cells represent the first targets of SARS-CoV-2 and are key sentinels in the initiation of early antiviral responses. To further determine viral tropism and heterogeneity of replication and antiviral responses in the presence and absence of remdesivir, we infected nHTBE cells with SARS-CoV-2 and analyzed single cell transcriptomes via the 10X Genomics platform. We captured an average of 1729 cells per experimental condition and 1836 variable genes per cell. We observed significant heterogeneity in the level of virus replication supported per cell (Fig. 2A, left). The frequency of cells with abundant viral RNA (defined as >0.1% of total reads per cell) closely matched what we observed using flow cytometry (Fig. 2A, right). Using this threshold for infection enables us to discriminate between cells that are productively infected from those with low-level viral RNA. Additionally, we confirmed viral replication kinetics with and without remdesivir by qRT-PCR, using a SARS-CoV-2 standard to calculate total viral genome copies (Fig. S1A). Together these results show a distinct impact of remdesivir treatment on SARS-CoV-2 viral reads per cell, consistent with our flow cytometry results.

**Fig. 2.**
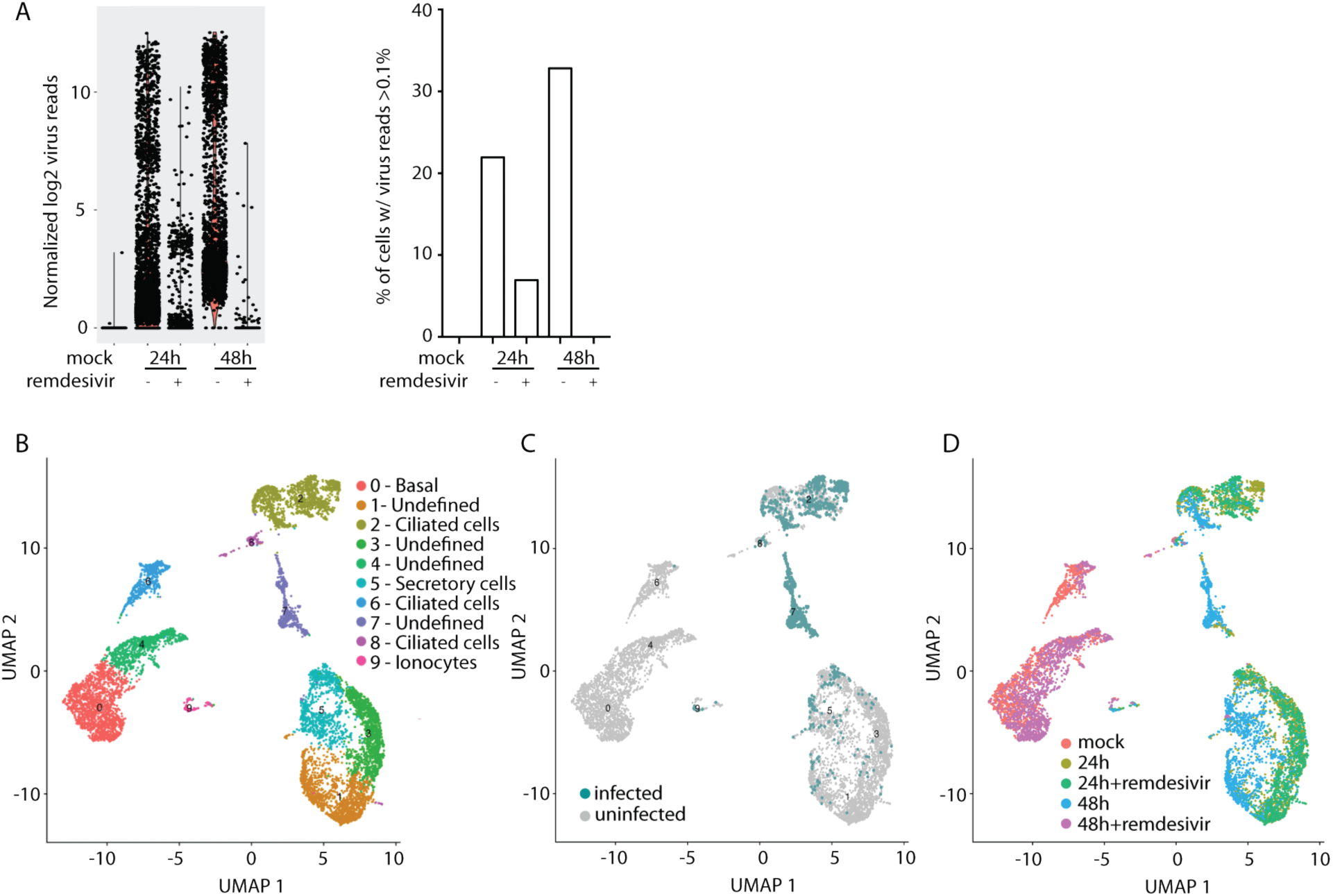
Single cell resolution of SARS-CoV-2 infection. (A) Total viral reads per cell in each condition (left) and frequency of cells expressing at least 0.01% virus reads (right). (B) Cells from all samples were combined and analyzed by UMAP. (C) Cells with greater than 0.1% of total reads mapping to SARS-CoV-2. (D) Cells colored according to experimental condition.

To determine SARS-CoV-2 tropism and the impact of remdesivir, we analyzed cells combined from all experimental conditions, including mock-infected and those at 24 or 48 hours post-infection (hpi), with or without remdesivir treatment, by uniform manifold approximation and projection (UMAP) [34, 35]. While some clusters of cells in this analysis were defined by cell type (basal, ciliated ionocytes), the majority were undefined (Fig. 2B and Supplemental table 2). We observed several clusters that were characterized by a preponderance of infected cells (Fig. 2C); each of these clusters contained cells from different experimental conditions (Fig. 2D). For example heavily infected cells in clusters 2 and 7 are derived from cells infected at 24hpi, 24hpi with remdesivir and 48hpi. Consistent with our flow cytometry results, cells of a variety of phenotypes localized to each of these dominant infected cell clusters. Interestingly, the majority of cells treated with remdesivir (Fig. 2A and Fig. S1A) clustered most closely with mock-infected cells. This clearly demonstrates the antiviral efficacy of remdesivir and suggests that the presence of active SARS-CoV-2 replication is the strongest driver of transcriptomic phenotype in this cell population. These data suggest that SARS-CoV-2 infected airway epithelial cells adopt diverse transcriptional profiles dominated by both viral RNA level and cell type.

### Single cell RNA sequencing identifies SARS-CoV-2 tropism in differentiated tracheal bronchial epithelial cells

Because UMAP clustering of *combined* samples was dominated by experimental condition, we next used t-squared neighbor embedded (t-SNE) clustering on *individual* samples to identify infected cell types and uninfected bystander cells (Fig. 3A-E and supplemental table 2). Ciliated cells were a major infected group at 24 and 48 hpi (Fig. 3B and D), consistent with our flow cytometry data. We also observed heavily infected cells that did not cluster by cell type at 48 hpi (Fig. 3D), possibly due to loss of cell markers during SARS-CoV-2 infection, as has also been observed with influenza infection [36]. Similar to what others have observed [11, 12, 37], we found low level expression of ACE2 in our primary human epithelial cells (Fig. 3A-E), likely reflecting the limited sensitivity of scRNA sequencing. We therefore validated these findings using qRT-PCR and detected low ACE2 (30.4 ct) which increased after IFNβ treatment (26.07 ct, ∼25 fold) (Fig. S1B), consistent with other studies [7, 33]. Interestingly, we also observed augmented of the host protease TMPRSS2, implicated in SARS-CoV-2 entry [32142651], within ciliated cells, secretory cells, and ionocytes, but reduced or no expression in basal cells (Fig. 3A-E). To determine if TMPRSS2 is the primary protease utilized for SARS-CoV-2 entry in differentiated nHTBE cells we treated cells with the TMPRSS2 inhibitor camostat. Camostat has been shown to block pseudovirus infection in undifferentiated primary human lung [5] and SARS-CoV-2 infection in immortalized Calu3 cells [5, 38]. Consistent with these results, blocking TMPRSS2 resulted in a near complete abrogation of infection in both ciliated and secretory cells (Fig. 3F). These data demonstrate that TMPRSS2 is the dominant protease entry factor for cells of the upper airway and may help explain the relative resistance of basal cells to infection with SARS-CoV-2. nHTBE cells pre-treated with remdesivir had reduced numbers of infected cells (Fig. 3C and E), similar to our flow cytometry data. Cell tropism of SARS-CoV-2 was not altered with remdesivir treatment, as infected cells were predominately ciliated cells and a few secretory cells at 24 hpi. Remdesivir pre-treatment had a stronger effect at 48 hpi as there were almost no infected cells detected.

**Fig. 3.**
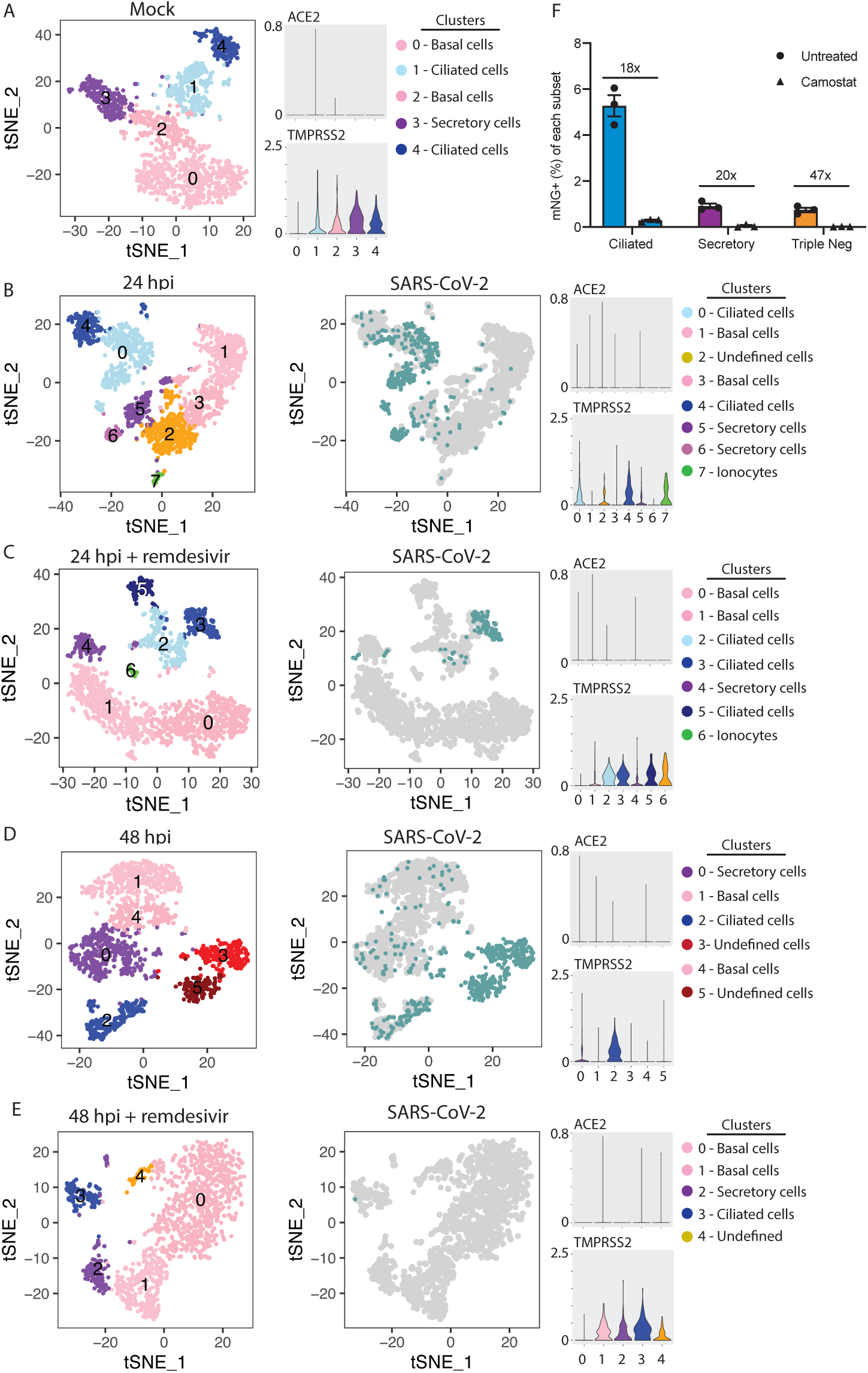
Single cell resolution of SARS-CoV-2 tropism. tSNE plots demonstrating clustering (left) and infected cells (right, blue) and volume plots for ACE2 and TMPRSS2 in mock (A), 24 hpi (B), 24 hpi with remdesivir (C), 48 hpi (D), and 48 hpi with remdesivir (E). (F) nHTBE cells were pretreated with vehicle or camostat, then infected with icSARS-CoV-2-mNG at MOI=2.5 and analyzed for mNG in ciliated (SiR-tubulin^+^ CD271^-^), secretory cells (CD66c^+^ SiR-tubulin^-^ CD271^-^), and triple negative cells. Numbers indicate fold change between vehicle and camostat treated cells.

### Early antiviral immune response to SARS-CoV-2 in human airway epithelium

To evaluate the relationship between SARS-CoV-2 infection and antiviral immune responses we measured the induction of type I and III IFNs among individual cells in the human airway epithelium. To determine if there is a relationship between the levels of virus and the induction of IFN, we plotted any cell expressing IFNB, IFNL1, IFNL2, IFNL3 or IFNL4 by the level of SARS-CoV-2 infection. There was a strong positive correlation between levels of SARS-CoV-2 replication and induction of IFN (Fig. 4A and B). Interestingly, we found cells with high levels of virus and low levels of IFN expression, potentially representing the successful evasion by SARS-CoV-2 of virus sensing pathways triggered by MDA5 or RIGI. At 48 hpi we observed two clusters of undefined cells, both with high SARS-CoV2 replication but divergent IFN responses: cluster 3 with high IFN induction and cluster 5 with reduced IFN. This disparate induction of IFN could indicate opposite outcomes of SARS-CoV-2 infection that drive differential transcriptomic clustering. We identified very few (∼0.5% at 24hpi) cells actively transcribing IFN, consistent with observations in other viral infections where a small number of IFN-producing cells is sufficient to establish the antiviral state [24, 25]. Despite this, we detected robust induction of ISGs (Fig. 4C). We evaluated ISGs with the greatest differences across sample condition and found several discrete patterns of expression. This analysis revealed ISGs that were only present in clusters of cells characterized with high levels of SARS-CoV-2 replication (Fig. 4C, MICB and GEM). We also observed an ISG (MT1F) that was preferentially induced within ciliated cells (Fig. 4C). At 48 hpi cells with the highest levels of viral RNA (clusters 3 and 5) had significantly lower levels of expression of several ISGs (DDIT, IFITM3, LY6E, TNFSF10), suggesting an impaired cellular antiviral immune response that enables unchecked SARS-CoV-2 replication (Fig. 4C).

**Fig. 4.**
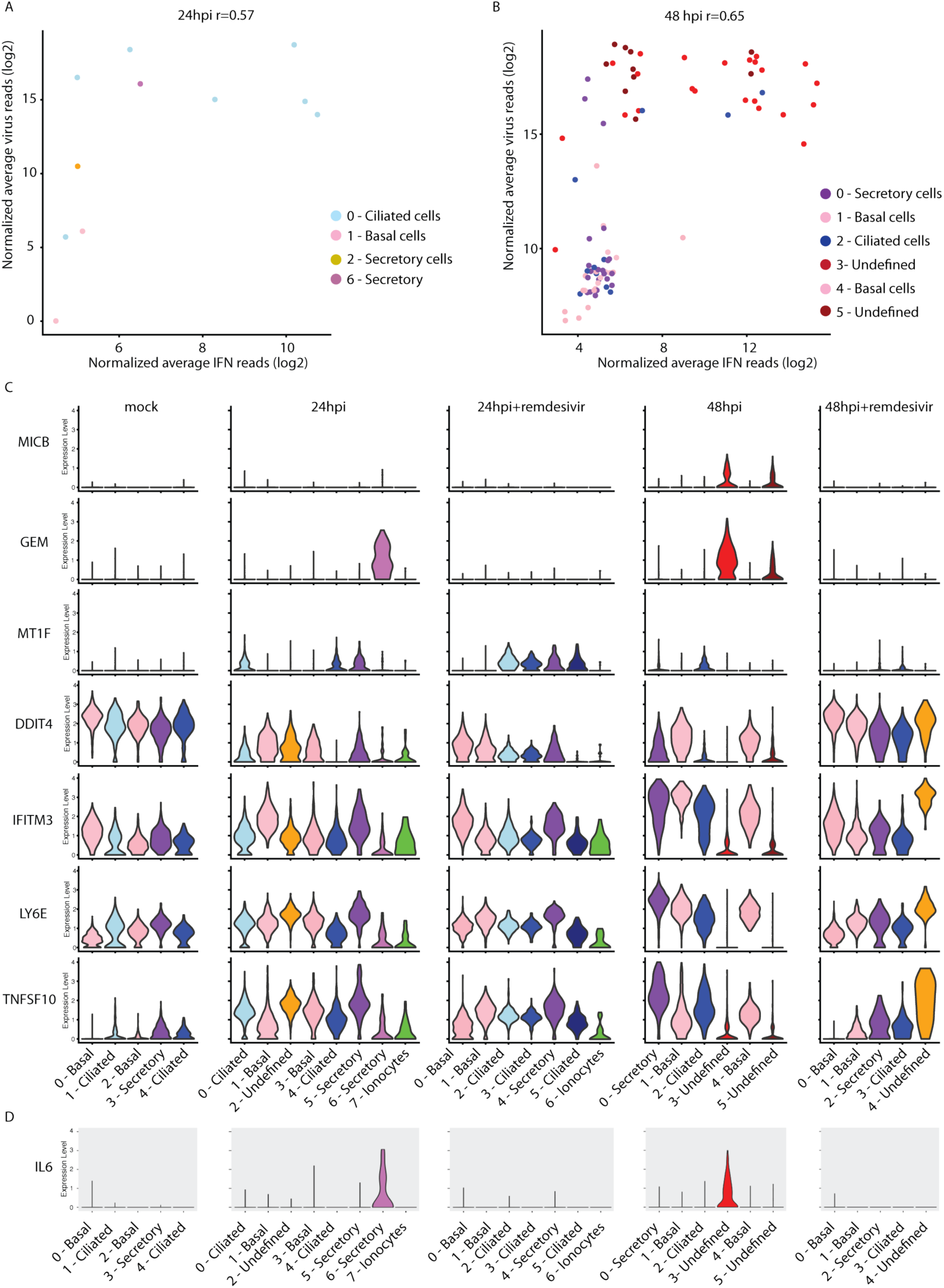
Distinct antiviral responses during SARS-CoV-2 infection. Cells from Fig. 3 were analyzed at 24hpi (A) and 48hpi (B) cells expressing interferon (IFNB1, IFNL1, IFNL2, IFNL3, IFNL4) were assessed for levels of combined IFN transcripts plotted against combined SARS-CoV-2 reads. Colors indicate clusters from Fig. 3B and D, respectively. (C) Volume plots of cells from Fig. 3 for ISGs with statistically significant expression between samples. (D) Volume plot of IL-6 expression.

Notably with SARS-CoV-2 infection we observed induction of IL-6, a pro-inflammatory cytokine implicated in the pathogenesis of COVID-19 [39], only within infected cells at 24 and 48 hours post-infection. Heavily infected secretory cells expressed the highest levels of IL-6 at 24 hpi (Fig. 4D). To determine the impact of early antiviral therapy on the hyperinflammatory response that drives COVID-19 pathology, we also analyzed infected cells pretreated with remdesivir. Remdesivir blocked the expression of IL-6 but not the induction of ISGs, suggesting that direct antiviral therapy can prevent induction of potentially pathogenic inflammation while preserving protective IFN responses.

## Discussion

The SARS-CoV-2 pandemic continues to spread, causing significant worldwide morbidity and mortality. Understanding early events in virus-host interactions in relevant model systems will be critical for understanding the pathophysiology of the disease as well as for evaluating antiviral and immunomodulatory drugs. Here we demonstrate that SARS-CoV-2 has preferential tropism for ciliated cells and secretory cells in the upper airway. Within these primary cells SARS-CoV-2 is only sensed by a small number of infected cells leading to IFN activation. Despite the low number of cells expressing IFN, there is a robust antiviral ISG response established. Finally, we demonstrate that remdesivir is capable of restricting virus replication uniformly across all susceptible cells within the upper airway.

A recent study has demonstrated increased accumulation of remdesivir triphosphate in human airway epithelial cultures compared to Calu-3 and Vero cell cultures [17]. This study only addressed the accumulation and efficacy in readouts from total airway epithelial cell cultures. Here we are able to observe remdesivir efficacy in all of our individually defined cell populations using actual viral replication as the readout. In addition to blocking virus replication, remdesivir may also provide benefits from blocking induction of proinflammatory cytokines while maintaining antiviral signaling. We uncovered differential expression of the entry factor TMPRSS2 across multiple cell types with higher expression in ciliated cells and near absent expression in basal cells. The protease inhibitor camostat blocked SARS-CoV-2 infection across all susceptible cells, suggesting TMPRSS2 is the dominant protease that enables infection in the upper airway. Differential expression of this and other host entry factors may explain the resistance of basal cells to infection.

Coronaviruses employ diverse mechanisms to evade detection and induction of IFN. The exact mechanisms SARS-CoV-2 uses are unclear. However, SARS-CoV-2 clearly deploys successful strategies as induction of IFN within infected cells is rare (Fig. 4A and B). While SARS-CoV-2 is proficient at evading IFN induction in most cells, it seems to be poor at countering effects of ISGs. Consistent with several recent reports, we demonstrate that SARS-CoV-2 is highly sensitive to IFN treatment (Fig. 1) [31-33]. We have previously identified ISGs in epithelial cells associated with high levels of virus replication [27]. Congruently, cells with high levels of SARS-CoV-2 reads had significantly increased expression of some of these same ISGs including GEM (Fig. 4D), RIPK2, PIM3, and ODC1 (Supplemental table 3). We have previously demonstrated a number of cell type specific ISGs during influenza A virus infection in mice [27]. Interestingly, we also identified a cell type-specific ISG (MT1F) induced during SARS-CoV-2 infection, which was primarily expressed in ciliated epithelial cells. A better understanding of both the heterogeneity of ISG responses and cell-type specific induction will help to determine transcriptional combinations that are successful in blunting infection versus those that lead to establishment of infection, robust replication, and spread to new cells.

Together the results presented here demonstrate that ciliated airway epithelial cells are the predominant cell type initially infected by SARS-CoV-2 and reveal heterogeneity within infected cells and in antiviral responses. Importantly, remdesivir is capable of reducing viral replication in all infected cell types within this culture system. We also discovered that after viral spread at 48 hpi, IFN responses and TMPRSS2 expression are the major determinants of virus tropism in susceptible cells. Our results demonstrate that infected epithelial cells at the earliest stages can produce IL-6, which potentially contributes to the cascade of inflammatory disease that occurs later during infection. Defining virus-host interactions at the single cell level reveals broad tropism and successful evasion of virus sensing, key attributes that likely allowed the zoonotic emergence of SARS-CoV-2.

## Materials and Methods

### Primary nHTBE cell culture

nHTBE cells (Lonza Bioscience, CC-2540S) harvested from healthy individuals were expanded using Pneumacult™-Ex Plus (Stem Cell Technologies) and seeded on 24 well hanging inserts, 0.4 µm pore (Corning, USA) and were maintained at ALI for 21^+^ days in a Pneumacult™ ALI growth medium (Stem Cell Technologies) at 37°C/5% CO_2_ in a humidified incubator. Cells were used 21^+^ days post ALI for experiments.

### Drug treatments and infection with SARS-CoV-2

Basal layer of medium was replaced with infection media (DMEM with 2% FBS, 100 U/mL penicillin and streptomycin, 10 mM HEPES, 0.1 mM nonessential amino acids, 4 mM L-glutamate, and 1mM sodium pyruvate). Wells were left untreated or treated in the basal compartment with 100 ng of IFNβ or 0.1 μM remdesivir for 1 hr prior to infection, or with 50 μM of camostat mesylate for 2 hrs prior to infection. For infection, SARS-CoV-2 strain 2019-nCoV/USA_WA1/2020 (provided by World Reference Center for Emerging Viruses and Arboviruses at the University of Texas Medical Branch) or icSARS-CoV-2-mNG, were added directly to the apical layer of cells for 1 hr at 37°C with rocking every 10 min. The virus was removed from the apical compartment and cells were incubated at 37°C until harvest.

### Flow cytometry analysis

Prior to harvest, cells were incubated in culture medium containing SiR-Tubulin (Spirochrome/Cytoskeleton Inc.) for 1 hr at 37°C. Cells were washed and trypsinized using ReagentPack™ Subculture Reagents (Lonza Bioscience) per the manufacturer’s instructions. Single cell suspensions were washed with 1X PBS and stained with Ghost Dye™ Violet 450 (Tonbo) for 30 min on ice. Cells were washed once with FACS buffer (cold HBSS supplemented with 2% bovine serum), stained with surface Abs, then fixed with 4% PFA in PBS, before flow cytometric detection on a BD LSR Fortessa (Becton Dickinson). Abs used include the following: CD66c (B6.2/CD66) (BD Biosciences); CD271 (ME20.4) (Biolegend); and TSPAN8 (458811) (Biotechne).

### Sample preparation for single-cell RNA sequencing

Cells were washed and trypsinized using ReagentPack™ Subculture Reagents (Lonza Bioscience). Single cell suspensions were washed with PBS + 5% FBS, resuspended in 200 µL PBS + 5% FBS, passed through a 70 µm filter, and placed on ice. Cells were counted using trypan blue exclusion on a Luna II (Logos Biosystems). The appropriate numbers of cells to achieve a targeted cell input of 5,000 cells per condition were used to generate the GEMs (Gel Bead-In Emulsion). The Chromium Next GEM Single Cell 3’ Gel beads v3.1 kit (10X Genomics, Pleasanton, CA, USA) was used to create GEMs following manufacturer’s instruction. All samples and reagents were prepared and loaded into the chip and ran in the Chromium Controller for GEM generation and barcoding. GEMs generated were used for cDNA synthesis and library preparation using the Chromium Single Cell 3’ Library Kit v3.1 (10X Genomics) following the manufacturer’s instruction.

### Single-cell RNA sequencing analysis

The 10x Chromium Single Cell sequencing raw data of experimental samples were reference mapped using Cell Ranger Count (version 3.0.1) for alignment to a hybrid human+SARS-CoV-2 viral genome reference (Homo_sapiens.GRCh38; MN985325). The hybrid genome reference, annotation file analysis scripts are available from our github site. The resulting raw count matrix for each experimental data set was imported into an R pipeline using Seurat 2.3.4 [40], where the filter criteria for empty droplets are minimum 1000 genes per cell, for genes that are presented in a minimum of three cells and for mitochondria genes percentage is no more than 30%. The default parameters were used for subsequent data normalization and scaling. The Seurat default method was used to select variable genes for PCA and clustering analysis, which the possible positive viral and mitochondria variable genes were excluded from final variable gene lists for all samples. The Seurat functions and utilities were used for subsequent cluster marker gene selection and results visualization. The ISG genes in the all marker list (FindAllMarkers function) of integrated analysis was ranked by their adjusted p value and ISGs within the top 30 that demonstrated disparate expression patterns are plotted with Seurat ViolinPlot function. To assess the relationship between all cells across all experimental conditions we used the FindClusters function with default parameters (except dim=1:16 and resolution = 0.15) and the RunUMAP function with default parameters (dims=1:16) of Seurat (version 3.2.2) was used for the integrated analysis for all five samples.

Overall cellular infection status (infected/uninfected) was determined by examining the distributions of percentage of viral counts within each cell on the log2 scale to magnify the differences at the low end. Thresholds for calling cells “infected” were set by calculating the percentage of summed viral counts within each cell over total counts, which is 0.1%. Cell types were defined using genes identified by Plasschaert et al [41]. A cutoff of at least 15 cell-type-specific genes was used to identify cell types for individual clusters.

### Quantitative RT-PCR

RNA was extracted from cells using RNeasy Plus Micro Kit (Qiagen) and cDNA was generated by reverse transcription using Superscript II (Invitrogen). Genes in each sample were amplified using gene-specific primers: Human_ACE2_F 5-CGAGTGGCTAATTTGAAACCAAGAA-3; Human_ACE2_R: 5-ATTGATACGGCTCCGGGACA-3. RNA was normalized to the endogenous alpha-tubulin primer probe set: 5’-GCCTGGACCACAAGTTTGAC-3’ and 5’-TGAAATTCTGGGAGCATGAC-3’. To determine the SARS-CoV-2 genome copy number in each sample, a standard curve was generated using serial dilutions of quantitative synthetic RNA (BEI #NR-52358) and virus specific nsp14 primers: 5’-TGGGGYTTTACRGGTAACCT-3’ and 5’-AACRCGCTTAACAAAGCACTC-3’. The genome copy number in each sample was derived from this curve.

## Data Availability

All sequencing data are available from NCI GEO accession number: GSE157526. All code used for single cell analysis, along with associated documentation, are available from: https://github.com/heznanda/scrnaseq-hybrid-cov2

## Acknowledgements

We thank the Minnesota Super Computing Institute, the World Reference Center for Emerging Viruses and Arboviruses and the University of Minnesota Medical School Deans office for support. This work was funded by R01 AI148669 to RAL, R01 AI153602 to VDM, T32 AI055433 to JMT, T32 HL007741 to WTM, and UL1TR002494 to SSS.

---

**Fig. S1.**
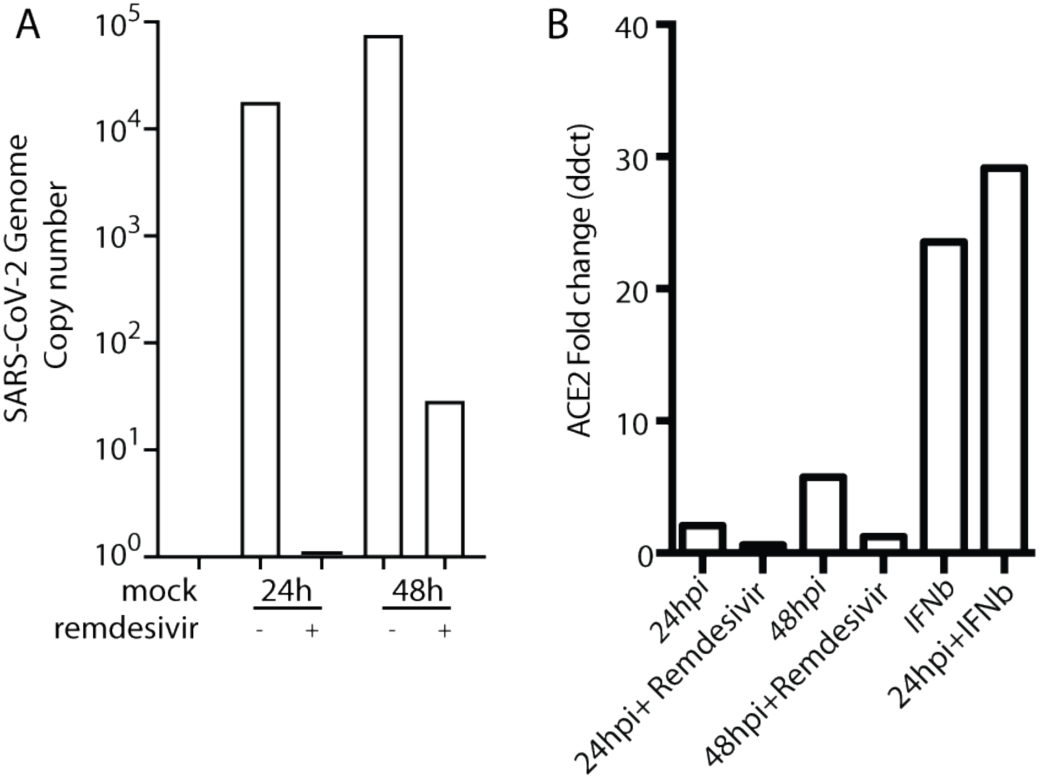
qRT-PCR validation of single cell RNAseq expression for SARS-CoV-2 and ACE2. Total RNA from cells used in Fig. 2 analyzed for (A) SARS-CoV-2 (log10(RNA copies per sample)) detected in cells samples collected at 24 or 48 hours after infection with SARS-CoV-2 in the presence or absence of remdesivir calculated by qRT-PCR. (B) Total RNA from cells in Fig. 2 and cells treated with or without IFNβ analyzed by qRT-PCR for ACE2.

